# RAWtunda: a Tool to Convert a Multi-Channel Raw Image into TIFF and OME-TIFF Formats

**DOI:** 10.1101/2025.10.25.684557

**Authors:** Natalia M. Dworak, Maxwell Cooper, Jay W. Fox, Ana K. de Oliveira

**Affiliations:** University of Virginia School of Medicine Spatial Biology Core, Charlottesville, VA, USA; Department of Microbiology, Immunology and Cancer Biology, University of Virginia School of Medicine, Charlottesville, VA, USA; Department of Pathology, University of Virginia School of Medicine, Charlottesville, VA, USA

**Keywords:** converter, MCD TIFF, OME-TIFF, ndpi, CosMx, IMC, digital pathology, spatial biology

## Abstract

**Motivation:** High-resolution biological imaging in spatial biology produces data in many proprietary formats. The lack of compatibility between these formats restricts reproducibility and analysis, creates access issues, and makes early data analysis a challenge.

**Results:** Here, we introduce a graphical user interface (GUI) application designed to convert proprietary image formats into standardized formats like OME-TIFF and TIFF which we have termed Rawtunda. This tool addresses the need for easy and efficient handling of large, complex imaging data generated in spatial biology and microscopy, facilitating data sharing, analysis, and long-term storage. Featuring an intuitive interface the application supports users in converting.mcd and.ndpi formats, generated from Image Mass Cytometry (IMC) and Digital Pathology scanners, respectively. This resource aims to improve interoperability of spatial biology datasets, streamline data management workflows, and promote reproducibility in imaging research and analysis, preserving crucial image metadata. The app ensures compatibility with downstream tools and is designed for both bioinformaticians and bench biologists without experience in coding.

**Availability and implementation:** The software, the documentation, and examples are available as open-source a https://med.virginia.edu/spatial-biology-core/rawtunda/ under the Copywrite of University of Virginia.

## Introduction

Advancements in biomedical sciences research technologies have dramatically increased the volume and complexity of its biological data. Spatial Transcriptomics (Marx 2021) and Proteomics (Nature Methods 2024) were considered the methods of the year in 2021 and 2024, respectively, and brings us the possibility to deeply explore the complexity of biological systems. It revolutionized our understanding of cellular and tissue architectures by enabling high-resolution imaging of biological specimens associated with information on analyte localization and expression.

Modern spatial biology instruments, such as those producing.mcd and.ndpi files, generate large and complex image datasets that are often stored in proprietary formats. While these formats are optimized for the specific imaging platforms, their closed nature poses significant challenges for data sharing, analysis, and long-term accessibility. Consequently, there is a growing need for tools that can efficiently convert proprietary image files into standardized, open formats such as Tag Image File Format (TIFF) and Open Microscopy Environment-TIFF (OME-TIFF), which are widely supported by most image analysis software and data repositories.

Each dataset typically consists of multi-channel, high-resolution images containing thousands to millions of cells, each with spatially resolved expression patterns. Analyzing such data requires computational pipelines that integrate image processing, cell segmentation, spatial statistics, and machine learning. For tasks such as: accurately segmenting cells, quantifying marker expression, identifying spatial domains, and modeling intercellular relationships, it is critical for transforming raw image data into meaningful biological insights without losing metadata information from the original raw file.

Proprietary formats of this data, specific to hardware vendors, makes integration with downstream analysis pipelines challenging. TIFF (Altheide 2011) and its extension, OME-TIFF (Goldberg 2005, Open Microscopy Environment 2023), have emerged as standards for biological image storage due to their support for high-resolution multi-dimensional data and metadata. However, converting existing datasets to these formats remains a barrier for many laboratories/cores/researchers.

Existing solutions for format conversion are often limited by their complexity and lack of user-friendly interfaces making them inaccessible to many researchers/users. For example, Windhager (2023) created *readimc*, an open-source Python package which enabled the extraction of information from.mcd files generated on imaging mass cytometry (IMC) (Chang 2017); another Python package is also available from Bodenmiller Group on Github (2021). While it is a powerful tool, it requires Python knowledge to process the raw file, which adds an extra challenge during data analysis.

To address these limitations, we developed a GUI-based application designed specifically for converting proprietary spatial biology and microscopy image formats into compatible, open standards. By providing an intuitive interface and robust processing capabilities, this tool aims to streamline the conversion and give capability to focus on further workflows, facilitate data sharing, and promote reproducibility in spatial biology research.

RAWtunda is freely available application, a cross-platform tool that simplifies the conversion of spatial biology/pathology/microscopy image files to TIFF and/or OME-TIFF while preserving spatial, temporal, and channel information as well as imaging metadata.

## Materials/Implementations

Currently Rawtunda supports conversions of the formats listed in Table 1.

**Table 1.**
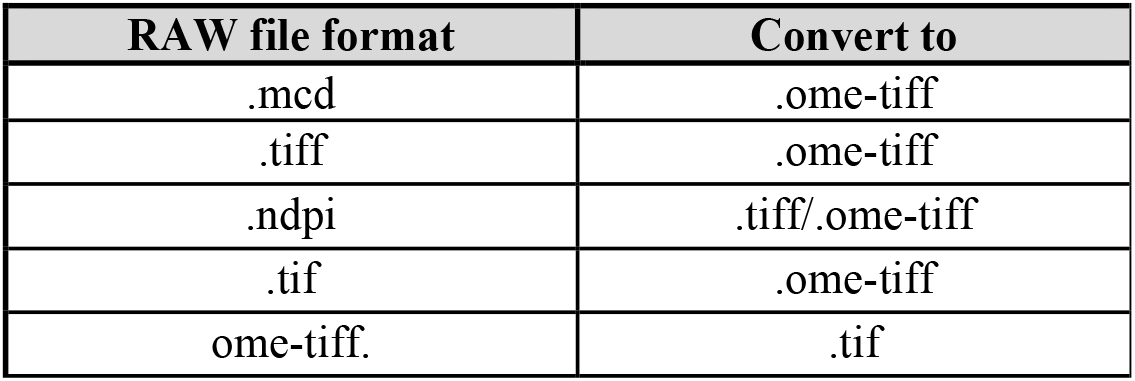
Overview of supported file conversions within the application, detailing input formats and target output formats.

A.ndpi file is a proprietary image file format developed by Hamamatsu for whole slide imaging, particularly in microscopy (https://openslide.org/formats/hamamatsu/). It is essentially a TIFF-like format with custom tags, designed to store large images, often gigabytes in size, from digital pathology scanners. While NDPI is a powerful format for storing high-resolution, multi-dimensional microscopy images, it requires specialized software for proper handling and conversion.

MCD files are a proprietary format for IMC analysis. Following image acquisition, the CyTOF mass spectrometry generates a single raw file for each region of interest (ROI) scanned, containing complete metadata information. Additionally, it produces a panorama image, which serves as the first screen of the entire tissue, akin to a bright field image without resolution. For example, region of interest (ROI) can be scanned in different resolutions such as tissue mode (5 μm resolution) or cell mode (1.0, 0.5, and 0.3 μm resolution). Further, the size of the mcd file increases with the resolution choice and tissue diameter, impacting the file conversion capacity due to the 4 GB limit of the TIFF file.

Our software conversion is limited to the size of TIFF (about 4GB). To work with files above 4GB it is necessary to work with BigTIFF format which requires a different conversion pipeline that does not lose metadata or distort the image. The update for this format is currently being tested and will be available soon.

### Converting vendor file format images via RAWtunda GUI

Spatial Biology instruments collect data into their own file formats that are not easily read and viewed. Most instruments vendor, have provided some form of software/viewer that allow access to view data or do simple adjustments. RAWtunda is a GUI tool to enable easy translation of these raw data to TIFF and OME-TIFF. Users will be able to use the data in the new format for further analysis.

### Example Application

Researchers/users will need to install RAWtunda on their computers prior to attempting this protocol.

Execute RAWtunda. The graphical user interface will appear as shown (Figure 1). On the left side of figure 1, you will see available function buttons:

**Figure 1.**
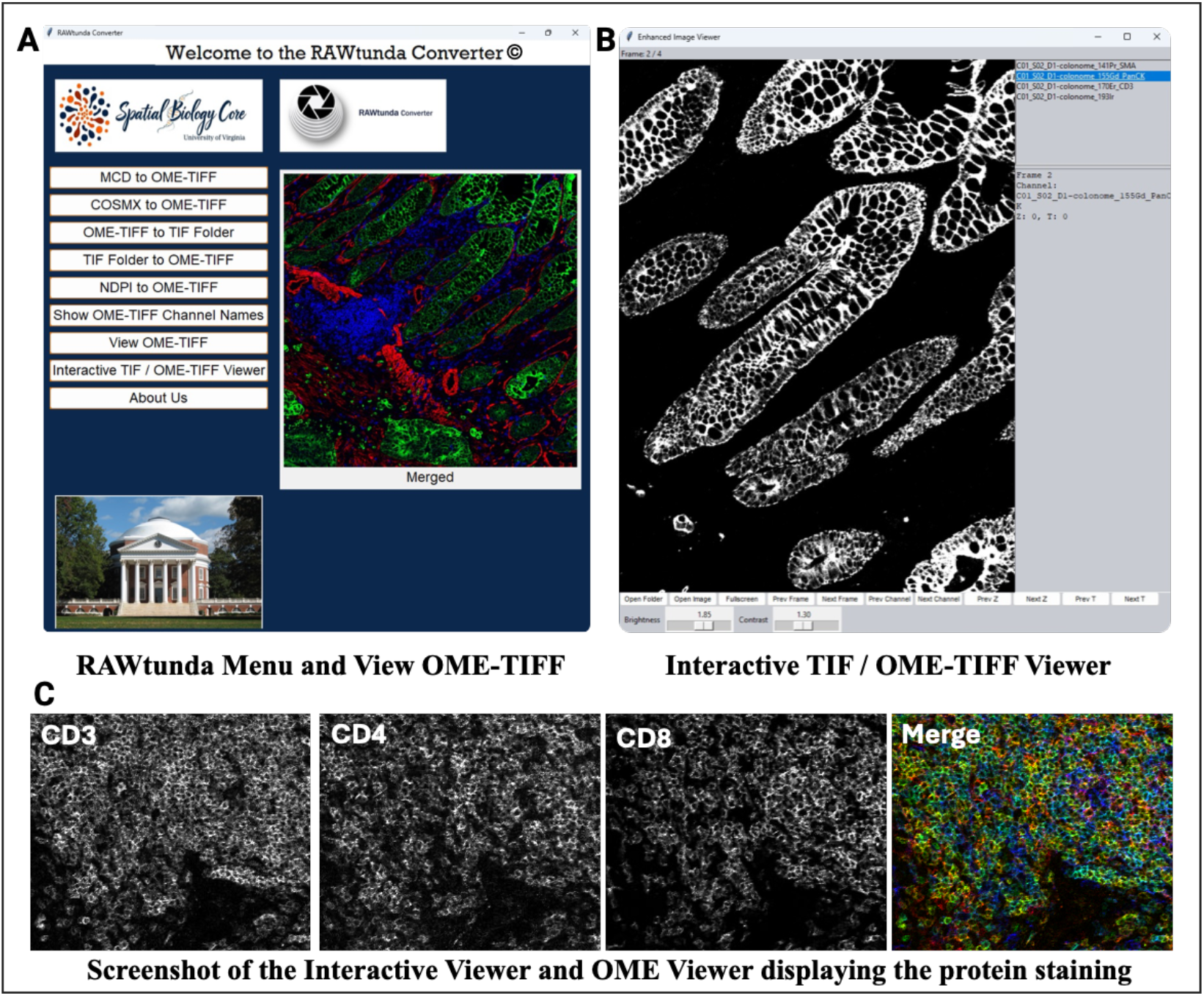
The main screen of the RAWtunda converter and viewer options. **(A)** The interface includes a primary navigation menu on the left, providing quick access to key functions from the home screen. The center area features a “View OME-TIFF” button for viewing colorized markers together. **(B)** On the right, all markers on the panel are displayed, allowing users to explore stain quality for each channel. The PanCK marker is highlighted in blue on the left and indicates epithelial cells in the colon cancer tissue in white in the center. **(C)** A panel shows different markers used to assess antibody staining quality: mouse spleen tissue stained for CD3 (green), CD4 (yellow), and CD8 (blue).

**MCD to OME-TIFF** – place your MCD File in the “Drop here” area or browse for file using “Browse for TIF File” button located underneath that area. The conversion will start automatically. The output OME-TIFF will have the same name as the input TIF file and will be located in the same folder. The software is designed to handle one ROI per round, only files containing one ROI can be converted.

**COSMX to OME-TIFF** - place your Cosmx File (single TIF File) in the “Drop here” area or browse for file using “Browse for TIF File” button located underneath that area. The output OME-TIFF will be located inside the input folder. The software converts one ROI per round.

**OME-TIFF to TIFF Folder** - Drop or select your OME-TIFF file.

The collection of newly created TIFs will be located in the same folder as the original OME-TIFF file. It is our recommendation that you place your OME in a folder of its own so that the created TIFs stay organized. The number of TIF images created is equal to the number of Channels in the original OME-TIFF file.

**TIF folder to OME-TIFF** - Drop or select the folder containing several TIF files to create one merged OME-TIFF file. Each TIF will be a single channel of the newly created OME-TIF file. Note, the end result is an OME-TIFF file located in the same location as the FOLDER that was selected. It is found under the folder, not inside the folder of the TIF Files. The created OME-TIFF will have the same name as the input folder.

**NDPI to OME-TIFF** - Drop or select your NDPI File to convert into an OME-TIFF File. Note that this conversion could take a while, due to a size of those files. A confirmation will appear when the conversion is complete.

The output could be a very large file because of how NDPI stores information. Ensure to only drop ONE file at a time.

**Show OME-TIFF Channel Names** - Drop or select an OME-TIFF file to view a list of its channel names. The entire functon of this button is being able to review the OME-TIFF channels.

**View OME-TIFF** - Drop or select an OME-TIFF file to see a ‘Merged’ view of all channels in the file. Before using this function, you can create an OME-TIFF file by selecting TIF files with specific markers (no more than 4) to generate a personalized image showing the co-localizations of your targets of interest. (Figure 1C and 1A)

**Interactive TIF/OME-TIFF Viewer** – displays a separate window, “Enhanced Image Viewer,” that provides extra features such as brightness and contrast adjustments and allows users to review each channel individually (Figure 1 B and 1C). This tool is particularly helpful for macOS users who previously could only access the mcd file image on a Windows system using MCD Viewer (Standard Bio Tools). Figure 1C displays a mouse spleen tissue image scanned with the Hyperion XTi in cell mode at 1 μm resolution. The interactive view mode enables users to explore each protein stain individually or together using View OME-TIFF.

## Results

### Compatibility

RAWtunda successfully converted files from.mcd (Hyperion XTi),.ndpi (Hamamatsu), and Nanostring/Bruker CosMx, formats to valid OME-TIFF or TIFF file formats and imported them into ImageJ, Fiji, or QuPath.

The metadata preservation enabled channel-specific view tracking and further analysis such as segmentation without manual correction.

The app was applied to.ndpi format images ranging between 1.4GB and 1.9GB and converted them into TIFF, keeping high resolution while zoomed in. The processing time differs depending on input file size.

### Operating Systems

The application is available in two versions:

- For Windows
- For macOS

The Windows version is optimized for Windows 10 and newer operating systems.

## Discussion

The proliferation of proprietary biological image formats has made standardization and interoperability a persistent challenge in image-based research. RAWtunda provides a practical and user-friendly solution by enabling reliable conversion of diverse image formats into TIFF and/or OME-TIFF, the latter being widely adopted, metadata-safe standard supported by major bioimage analysis tools.

The development of a GUI-based application capable of converting proprietary spatial biology/microscopy image formats into standardized open formats addressed a critical need within the research community. By supporting and enabling conversion to OME-TIFF and TIFF, the tool enhances data accessibility and interoperability. Preservation of essential metadata ensures that researchers can efficiently manage datasets without compromising data integrity.

Furthermore, user experience is central to the utility of this tool. The GUI design aims to lower technical barriers, making format conversion accessible to researchers without extensive computational expertise. Feedback from initial users indicates that ease of use and reliability are key factors for successful adoption. By bridging the gap between proprietary formats and open standards, RAWtunda empowers individual scientists without bioinformatics background to create reproducible and portable imaging workflows.

## Conclusion

This application represents a step forward in simplifying the management of biology imaging data. Continued development and community feedback will be essential to address emerging needs.

By the use of GUI, the tool is accessible to a broad range of users, from researchers with minimal programming experience to computational researchers integrating it into automated pipelines.

Future improvements will focus on expanding support for additional proprietary formats, and optimizing processing speed for extremely large datasets. Going forward we aim to extend support in analysis pipelines and offer more option for converting wider range of file formats. Ultimately, RAWtunda contributes to the growing ecosystem of open-source bioimage tools that promote reproducibility, transparency, and FAIR (Findable, Accessible, Interoperable, Reusable) principles in the life sciences.

## Availability

RAWtunda is available on https://med.virginia.edu/spatial-biology-core/rawtunda/ Copyright of University of Virginia (UVA), 2025.

## Acknowledgments

We acknowledge the use of the University of Virginia School of Medicine Spatial Biology Core Facility, RRID: SCR_023281; Flow Cytometry Core Facility, RRID :SCR_017829, and Research Histology Core, RRID: SCR_025470; UVA Comprehensive Cancer Center and the UVA Undergraduate Scholars Program.

